# Mitochondrial DNA metabolism is coupled with 20S proteasome function via regulation of deoxyribonucleotide homeostasis in *Saccharomyces cerevisiae*

**DOI:** 10.1101/2021.11.30.470679

**Authors:** Xiaowen Wang, Arnav Rana, Liam P. Coyne, Daniel M. Loh, Thuy La, Xin Jie Chen

## Abstract

The synthesis of mitochondrial DNA (mtDNA) is not coupled with cell cycle. Previous studies have shown that the size of deoxyribonucleoside triphosphate (dNTP) pools plays an important role in regulating mtDNA replication and amplification. In yeast, dNTPs are synthesized by the cytosolic ribonucleotide reductase (RNR). It is currently poorly understood as to how RNR activity is regulated in non-dividing or quiescent cells to finely tune mtDNA metabolism to cope with different metabolic states. Here, we show that defect in the 20S proteasome drastically destabilizes mtDNA. The mtDNA instability phenotype in 20S proteasome mutants is suppressed by overexpression of *RNR3* or by the deletion of *SML1*, encoding a minor catalytic subunit and an intrinsic inhibitor of RNR respectively. We found that Sml1 is stabilized in the 20S proteasomal mutants, suggesting that 20S affects mtDNA stability by stabilizing Sml1. Interestingly, defect in the regulatory 19S proteasomal function has only subtle effect on mtDNA stability, supporting a role of the 20S proteasome in dNTP homeostasis independent of 19S. Finally, we found that when cells are transitioned from glycolytic to oxidative growth, Sml1 level is reduced in a 20S-dependent manner. In summary, our study establishes a link between cellular proteostasis and mtDNA metabolism through the regulation of dNTP homeostasis. We propose that increased degradation of Sml1 by the 20S proteasome under respiratory conditions provides a mechanism to stimulate dNTP synthesis and promote mtDNA amplification.

## INTRODUCTION

Mitochondrial DNA (mtDNA) encodes a subset of proteins that directly participate in oxidative phosphorylation (OXPHOS), and ribosomal RNAs and tRNAs that are required for the synthesis of the mtDNA-encoded OXPHOS components inside mitochondria. Loss of mtDNA integrity directly affects cell’s energy metabolism. This leads to cell death due to energy deficiency and defects in other mitochondria-associated functions including stress signalling, iron sulfur cluster biosynthesis, cellular proteostasis and programmed cell death. The importance of understanding the mechanism of mtDNA maintenance is underscored by many human diseases that result from mtDNA mutation, deletion, and depletion. These clinical conditions can be caused by direct lesions to mtDNA, due to replicative errors and constant damage by reactive oxygen species. They can also be caused by mutations in nuclear genes that affects mtDNA replication and integrity (Wallace 2005; Copeland 2008; Vafai and Mootha 2012; Suomalainen and Battersby 2017). These genes include those that directly participate in mtDNA replication, such as *POLG1* and *POLG2* encoding subunits of the mitochondrial DNA polymerase (or Polγ), and *TWNK* that encodes the replicative Twinkle helicase.

Mutations in several other genes have been identified that cause mtDNA diseases by affecting the homeostasis of deoxyribonucleotide pools, including *RRM2B* that encodes the p53R2 subunit of ribonucleotide reductase (RNR). RNR is involved in reducing ribonucleoside diphosphates to deoxyribonucleoside diphosphates in the cytosol. The resulting deoxyribonucleotides are then imported into mitochondria, providing precursors for the generation of deoxyribonucleoside triphosphates (dNTPs) that support mtDNA replication. RNR is a tetrameric protein, made up of two dimeric subunits: a large catalytic subunit, R1, and the smaller R2 subunit. In mammalian cells, R1 and R2 move to the nucleus during S phase to promote dNTP synthesis and DNA replication. *RRM2B* encodes another R2 protein (or p53R2) and mutations in the gene lead to severe mtDNA depletion and multiple deletions (Bourdon *et al*. 2007; Tyynismaa *et al*. 2009). It has been shown that Rrm2B resides in the cytosol with a primary function in supporting mtDNA replication (Pontarin *et al*. 2008).

Multiple deletions in mtDNA are also associated with diseases affected in several mitochondrial inner membrane (IMM) proteins. For example, mutations in Mpv17 cause mtDNA-depletion diseases (Calvo *et al*. 2006; Spinazzola *et al*. 2006). A recent study showed that there is a significant reduction of dNTPs in mitochondria lacking Mpv17 (Dalla rosa *et al*. 2016), which further highlights the importance of dNTP homeostatsis for the maintenance of mtDNA integrity. In addition to *MPV17*, missense mutations in *ANT1*, involved in ATP/ADP exchange across the IMM, have also been found to cause dominant diseases autosomal dominant Progressive ophthalmoplegia (adPEo) manifested by deletions and depletion of mtDNA (Kaukonen *et al*. 1999; Kaukonen *et al*. 2000; Thompson *et al*. 2016). However, the underlying mechanism of mtDNA instability is much less understood. The mutant variants of Ant1 often have moderately reduced nucleotide transport activities (Fontanesi *et al*. 2004). Recent studies suggested that the dominant pathogenic variants of Ant1 and its equivalent in yeast, Aac2, clog mitochondrial protein import (Coyne et al., 2021, manuscript under review). This raises the possibility that mtDNA instability may result from a defect in the import of proteins critical for dNTP uptake and/or other mtDNA metabolism processes.

The synthesis of dNTPs mainly occurs during the S phase of cell cycle and is normally kept at a very low level in non-dividing or quiescent cells. As mtDNA replication is not coupled with cell cycle, a fundamental but unsolved question is how dNTP synthesis is regulated in non-dividing cells to support mtDNA replication and repair especially when cells need to amplify mtDNA for supporting oxidative metabolism. In this report, we show that reduced 20S proteasomal function in yeast greatly compromises mtDNA stability by affecting the level of Sml1, an intrinsic inhibitor of RNR. The data support the idea that the 20S proteasome plays an important role in coupling mtDNA replication with respiratory growth through the regulation of RNR activity and dNTP homeostasis.

## MATERIALS AND METHODS

### Growth media and construction of yeast strains

Yeast complete medium (YP) containing 1% Bacto yeast extract and 2% Bacto peptone is supplemented with 2% glucose (YPD) or 2% galactose (YPGal) or 2% glycerol (YPGly). Yeast minimal medium (YNBD) contains 0.67% Difco yeast nitrogen base without amino acids and 2% glucose. Supplements essential for auxotrophic strains were added to 20 μg/ml for bases and amino acids except for leucine (30 μg/ml). For selecting G418^R^ transformants, geneticin was added to YPD at 300 μg/ml. Unless specified, all the yeast strains used in this study are isogenic to M2915-6A or M2915-6alpha. Most null alleles of genes marked by *kan* were transferred from strains of the yeast knockout collection into the M2915-6A background by PCR amplification and one-step gene replacement. *SML1* was disrupted by the insertion of *URA3*. 3xHA tag was added to the C-terminus of Irc25, Poc4 and Pre9 by integration of a PCR product amplifying from a 3HA-KanMX6 cassette.

### Western blot and antibodies

Cell extracts were prepared as previously described (Chen 2001) and analyzed by SDS-PAGE followed by western blot using antisera against Sml1 (courtesy from Rodney Rothstein laboratory, Columbia University) or HA (Covance#MMS-101R). Total proteins were stained with REVERT Total Protein Stain (#926-11011, LI-COR). For western blot, band intensities were quantified by the LI-COR Odyssey Fc Imaging system and normalized against total protein staining or the level of the cytosolic Pgk1 protein.

### Petite frequency measurements

All the yeast strains used in this study have the *ade2* auxotrophic marker so that petites can be identified as white colonies on YPD. Cells were first grown on YPG plates for two days at 30°C and then inoculated into 2 ml of YPD broth at OD600 about 0.01. Cells were grown at 25, 30 or 37°C for 24 hours. The cultures were then diluted into fresh YPD medium at 1:1000 every 24 hours. At each time point, cells were diluted in water, plated on YPD medium and incubated at 30°C for 3-5 days. Petites were scored as white colonies and totally 200 −500 colonies were counted for each strain.

### Multicopy suppressor screen

A 2 μm-based yeast multicopy genomic library was transformed into the *poc4Δ* mutant. Approximately 5,000 Leu2^+^ transformants were manually picked up, diluted into water in a 96-well microplate, and spotted onto YPD plates. The cells were incubated at 37°C for five days. Red colonies were scored. Yeast plasmids were rescued from three red colonies, amplified in *Escherichia coli* and analyzed by Sanger sequencing. Two clones contain the *POC4* gene and the third clone has only the *RNR3* gene as a full length ORF in the insert DNA.

### Other techniques

Mitochondria were isolated as described by Boldogh and Pon (Boldogh and Pon 2007). Briefly, yeast cells were grown in YPGal medium, washed with Tris-SO4-DTT, treated with 5 mg zymolyase per gram of yeast to create spheroplasts and then broken with a 40 ml Dounce homogenizer. Mitochondria were collected by differential centrifugation. BN-PAGE was performed using the Life Techonologies NativePAGE™ Novex^®^ Bis-Tris Gel System as recommended by the manufacturer. Visualization of F1Fo-ATP synthase complexes by in gel ATPase activity staining was performed according to a previous established procedure (Bornhovd *et al*. 2006).

## RESULTS

### Defect in 20S proteasomal function destabilizes mtDNA

The proteasome is composed of heteroheptameric α-subunit and β-subunit rings that form the catalytic 20S proteasome, and the 19S regulatory particle (Hochstrasser 1996). Specific chaperones are required for the assembly of the 20S proteasome and the 19S regulatory particle. The yeast *AAC2* gene encodes the major isoform of ATP/ADP translocase on the inner mitochondrial protein. In a previous study, we found that overexpression of *POC4* and *UMP1*, two chaperone proteins for the assembly of the 20S proteasome (Ramos *et al*. 1998; Le tallec *et al*. 2007; Kusmierczyk *et al*. 2008), suppress cell death induced by *aac2^A128P^* (Wang and Chen 2015), equivalent to human *ant1^A114P^* that causes adPEO manifested by multiple mtDNA deletions. Interestingly, we found that overexpression of these two genes also supress the formation of respiratory deficient petite colonies in a strain expressing *aac2^M114P^* (Fig. 1A). M114P is equivalent to the L98P mutation in human *ANT1* that also causes adPEO and multiple mtDNA deletions (Napoli *et al*. 2001). To understand the link between proteasomal function and mtDNA stability, we asked whether disruption of these genes destabilizes mtDNA. Indeed, we found that mtDNA becomes highly unstable in yeast strains disrupted of *POC4* and *UMP1* when cells were grown in YPD medium at 25°C and 30°C (Fig. 1B and 1C).

**Figure 1.**
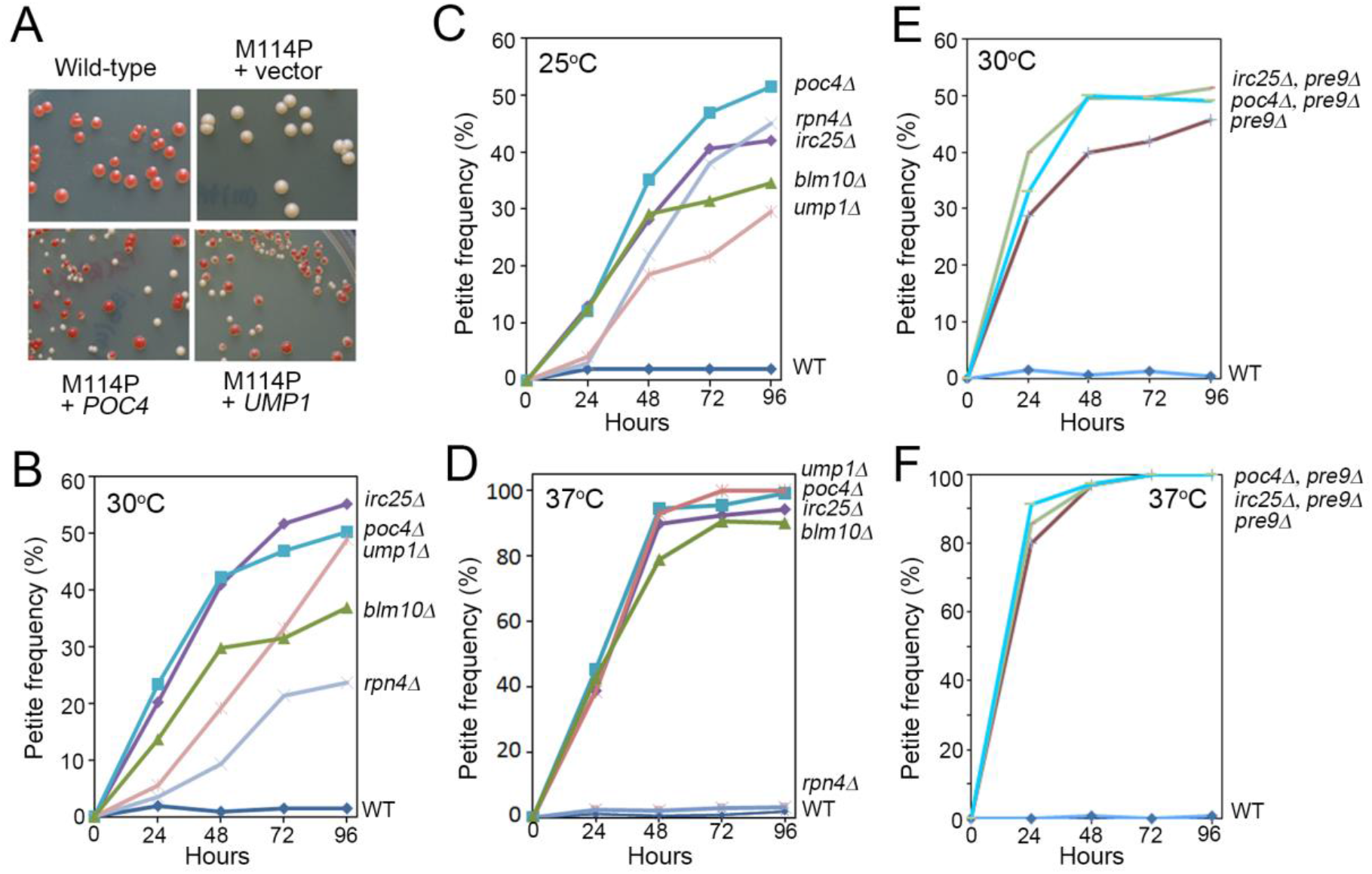
Overexpression and disruption of 20S proteasomal genes affect mtDNA stability. (A) Overexpression of *POC4* and *UMP1* from a multicopy vector suppresses the production of petite (white) colonies in cells expressing the chromosomally integrated *aac2^M114P^* (M114P) allele. The strain CS1460-2D was transformed with the plasmids expressing *POC4* and *UMP1*. Leu^+^ transformants were diluted in water and plated on YPD medium to allow colony formation at 30°C for 4-5 days. (B) – (F) Petite frequency in 20S proteasomal mutants grown in liquid YPD medium at temperatures as indicated.

To determine whether mtDNA instability results from a general defect in 20S proteasomal function, we generated strains lacking *IRC25* (or *POC3*) and *BLM10. IRC25* cooperates with *POC4* to orchestrate the assembly of the α3 subunit into the 20S proteasome. *BLM10*, the homolog of the human PA28 protein, is an activator of the 20S proteasome (Schmidt *et al*. 2005). We found that the mutant strains lacking these genes produce petite colonies at both 25°C and 30°C (Fig. 1B – 1C). We also found that disruption of *RPN4*, encoding a transcriptional factor that activates the expression of proteasomal genes (Mannhaupt *et al*. 1999; Xie and Varshavsky 2001), induces petite production at 25°C and 30°C (Fig. 1B-1C). The data suggest that the 20S proteasomal function is required for maintaining mtDNA stability.

To directly demonstrate a role of the 20S proteasome in maintaining mtDNA stability, we disrupted *PRE9* that encodes the α3 subunit of 20S. *PRE9* is the only nonessential 20S gene, as α3 can be replaced by the α4 subunit to generate an alternative proteasome with an additional copy of α4 (Velichutina *et al*. 2004; Kusmierczyk *et al*. 2008). As expected, we found that disruption of *PRE9* induces petite production (Fig. 1E). The α3 subunit is therefore important for mtDNA stability, which cannot be replaced by the α4-α4 proteasome. The *pre9Δ irc25Δ* and *pre9Δ poc4Δ* double mutants only slightly increase petite production compared with the *pre9Δ* single mutant (Fig. 1E), supporting the idea that disruption of *IRC25* and *POC4* destabilizes mtDNA stability mainly due to the loss of the α3 subunit.

### Global proteostasis in the cytosol affects mtDNA stability

One possible explanation for reduced mtDNA stability in 20S proteasomal mutants is that low proteasomal activity directly reduces mitochondrial quality by affecting processes such as mitochondrial dynamics, mitophagy and degradation by protein retro-translocation (Bragoszewski *et al*. 2017). mtDNA instability has previously been reported in the *blm10* mutant that affects mitochondrial fission (Tar *et al*. 2014). To better understand how proteasomal function affects mtDNA stability, we first examined whether loss of *IRC25* and *POC4* results in mitochondrial function that secondarily affects mtDNA stability. The *irc25Δ* and *poc4Δ* mutants have reduced growth on the non-fermentable glycerol medium, which is consistent with increased mtDNA instability (Fig. 2A). Using blue-native polyacrylamide gel electrophoresis (BN-PAGE), we examined the assembly of respiratory supercomplexes. We found that the assembly of the respiratory supercomplexes is little affected in the cells lacking *IRC25* and *POC4* (Supplemental Fig. 1). Consistent with this, in-gel activity staining showed that the assembly of the dimeric form of the complex V (V_dim_), which plays a critical role in morphological control of mitochondrial cristae (Paumard *et al*. 2002), is little affected by the loss of *IRC25* and *POC4* (Fig. 2B), despite limited release of the free F1 complex due to the loss mtDNA and the membrane Fo sector of the enzyme. Like in *blm10* mutant which increases mitochondrial fission (Tar *et al*. 2014), we also found that mitochondria are more fragmented in *irc25Δ* and *poc4Δ* mutants compared with wild type (Fig. 2C).

**Figure 2.**
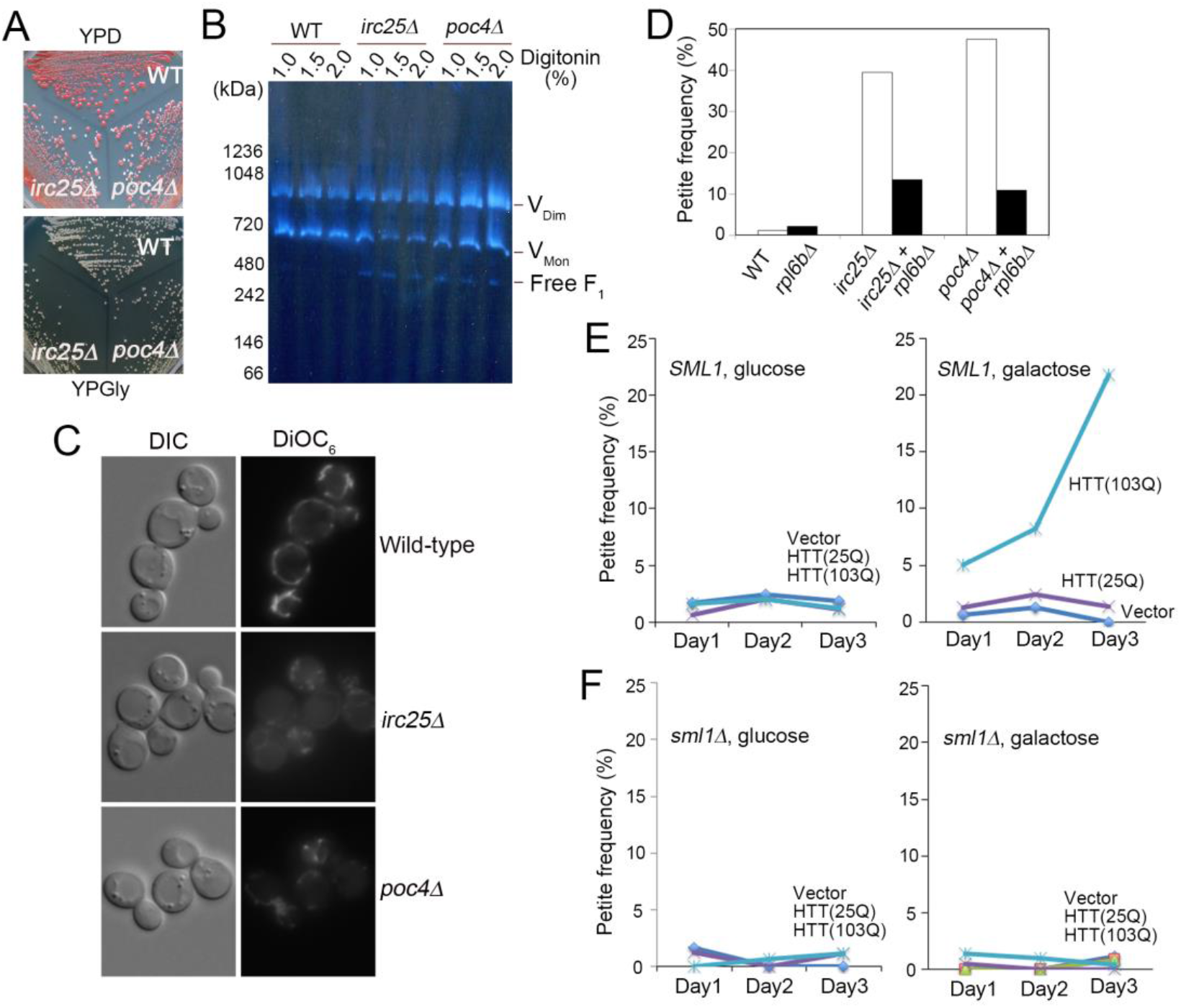
Global proteostatic stress affects mtDNA stability. (A) Respiratory growth on YPGly medium is moderately reduced in *irc25Δ* and *poc4Δ* mutants. (B) In-gel activity staining showing that the biogenesis of the monomeric (V_Mon_) and dimeric (V_Dim_) forms of F_1_F_o_-ATP synthase is little affected in the *irc25Δ* and *poc4Δ* mutants. (C) Mitochondrial fragmentation in *irc25Δ* and *poc4Δ* mutants. (D) Petite production in the *irc25Δ* and *poc4Δ* mutants is suppressed by the disruption of *RPL6B* that reduces global protein synthesis. (E) Induction of petite colonies in wild-type yeast cells expressing human HTT(103Q) but not HTT(25Q). Transformants of plasmids expressing GAL10-controlled HTT(103Q) and HTT(25Q) were grown in selective medium for 1-3 days as indicated before being plated onto YPD medium for scoring petite colonies. (F) Disruption of *SML1* suppresses petite production from HTT(103Q)-expressing cells.

We subsequently found that global cytosolic proteostasis affects mtDNA stability First, production of petites from all the 20S mutants with the exception of *rpn4Δ* is drastically increased when cells were incubated at 37°C (Fig. 1D and 1F). Secondly, disruption of *RPL6B*, encoding a cytosolic ribosomal protein, suppressed petite production in *irc25Δ* and *poc4Δ* cells (Fig. 2D). Disruption of *RPL6B* only mildly reduces protein synthesis and cell growth (Wang *et al*. 2008). This appears to be sufficient to improve mtDNA stability in *irc25Δ* and *poc4Δ* mutants. The data suggest that loss of mtDNA stability in the *irc25Δ* and *poc4Δ* mutants is likely caused by a defect in the degradation of a cytosolic protein. Disruption of *RPL6B* likely stimulates the process by improving cytosolic proteostasis and increasing the availability of proteasome for protein turnover. Thirdly, we found that protein misfolding in the cytosol destabilizes mtDNA (Fig. 2E and 2F). We challenged the cytosolic proteostasis of yeast cells by expressing the human huntingtin proteins from the galactose-inducible *GAL10* promoter (Meriin *et al*. 2002). We observed an increase in petite production when the aggregation-prone Htt103Q variant, which contains an expanded polyglutamine stretch, was expressed. No increased petite production was observed in cells expressing the non-aggregating Htt25Q variants. The data further support the idea that the proteostatic state of the cytosol affects mtDNA stability.

### Overexpression of *RNR3* and disruption of *SML1* suppress mtDNA loss in 20S mutants

To understand how mutations in 20S proteasomal genes reduce mtDNA stability, we performed a multicopy suppressor screen for genes that suppress mtDNA instability in the *poc4Δ* mutant. The *poc4Δ* cells in the *ade2* background form only white colonies at 37°C due to the loss of mtDNA. After transformation with a genomic library based on a multicopy vector, we examined ~5,000 transformants individually for their ability to form red colonies indicative of mtDNA retention when grown at 37°C on complete glucose medium (YPD). Three suppressor clones (SOP, for Suppressor of *poc4Δ*) were identified (Fig. 3A). Sequence analysis of the suppressor clones showed that the insert DNA in the SOP3 and SOP4 clones contains the *POC4* gene, whereas SOP16 contains the *RNR3* gene. *RNR3* encodes a minor isoform of the large subunit of RNR that catalyzes the rate-limiting step of dNTP biosynthesis. RNR is a tetrameric protein complex, consisting of two large (Rnr1 and Rnr3) and two small (Rnr3 and Rnr4) subunits in yeast (Fig. 3B). We found that overexpression of *RNR1*, encoding the major isoform of the large subunit of RNR, also suppresses the loss of mtDNA from *poc4Δ* cells (not illustrated). The data suggest that defect in 20S proteasomal function may reduce RNR activity and dNTP biosynthesis, which in turn affects mtDNA stability. Supporting this, we found that the *irc25Δ* and poc4Δ mutants are hypersensitive to hydroxyurea (HU), an inhibitor of RNR (Lammers and Follmann 1984).

**Figure 3.**
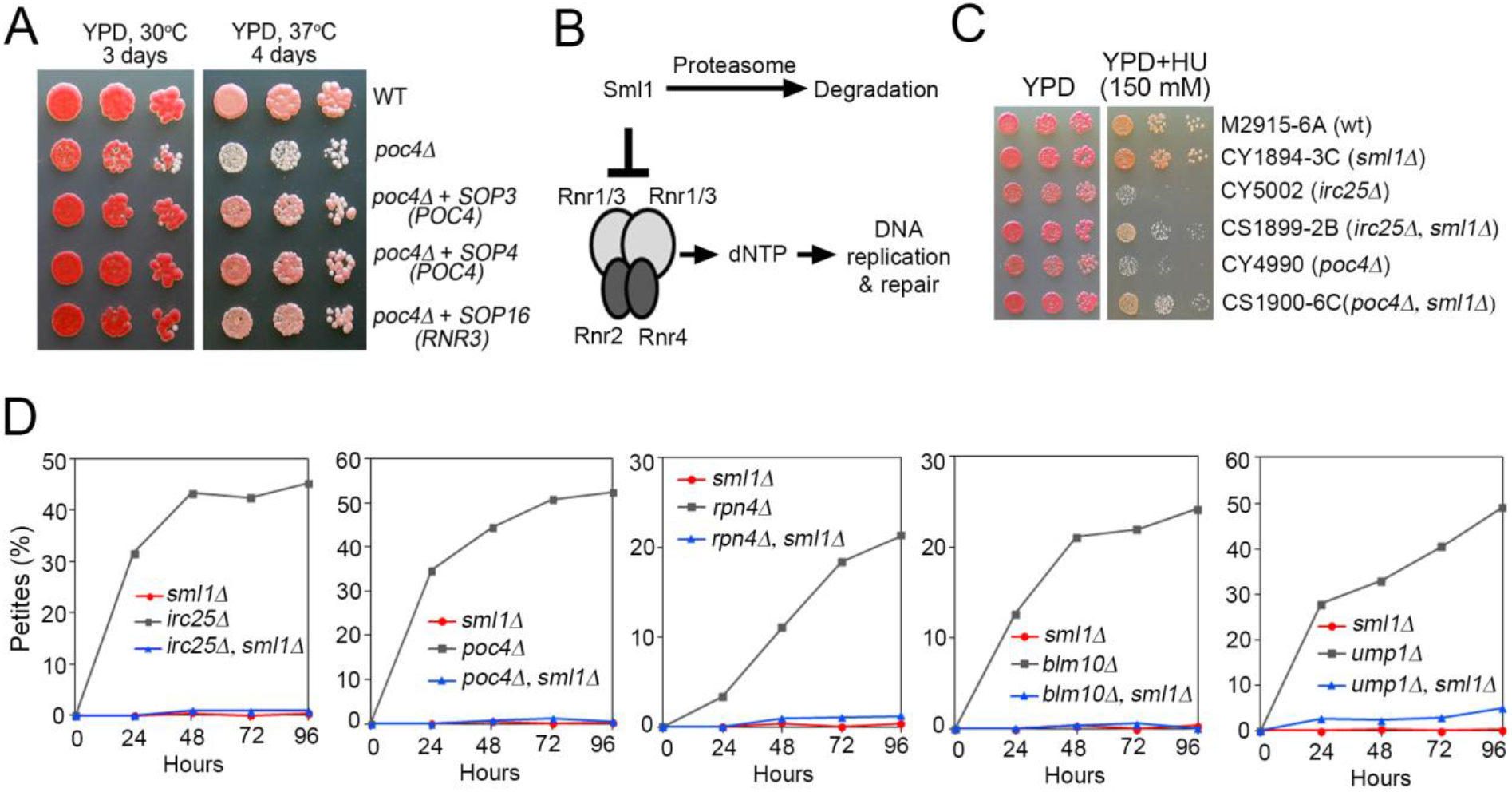
Genetic interaction between 20S proteasome mutants and the RNR pathway. (A) Overexpression of *RNR3* suppresses the formation of white (petite) colonies in the *poc4Δ* mutant on YPD medium. (B) Schematics depicting the inhibition of RNR by Sml1. (C) Sensitivity of *irc25Δ* and *poc4Δ* mutants to hydroxyurea, which is partially suppressed by the disruption of *SML1*. (D) Disruption of *SML1* suppresses petite production from 20S proteasome mutants.

RNR is regulated at multiple levels, including transcriptional activation, nuclear translocation, allosteric control and the direct inhibition of enzymatic activity by the Sml1 protein (Zhao *et al*. 1998; Chabes *et al*. 1999; Zhao *et al*. 2000; Zhao *et al*. 2001) (Fig. 3B). We found that disruption of *SML1* suppresses the hypersensitivity of the *irc25Δ* and poc4Δ mutants to HU (Fig. 3C), consistent with the idea that loss of Irc25 and Poc4 reduces RNR activity. More importantly, we found that deletion of *SML1* robustly suppresses the mtDNA instability phenotype in cells disrupted of *IRC25, POC4, RPN4, BLM10* and *UMP1*. Furthermore, mtDNA instability in cells expressing HTT(103Q) is also suppressed by the disruption of *SML1* (Figure 2F). These data strongly suggest that defect in 20S activity and global proteostatic stress destabilize mtDNA by reduced RNR activity.

### Stabilization of Sml1 in 20S proteasomal mutants

Sml1 is a highly unstable protein and is degraded in response to DNA damage which relieves RNR from inhibition and activates dNTP synthesis (Zhao *et al*. 1998; Chabes *et al*. 1999; Zhao *et al*. 2000; Zhao *et al*. 2001). We hypothesized that the mtDNA instability phenotype in 20S proteasomal mutants is caused by reduced turnover of Sml1, which in turn inhibits RNR and dNTP synthesis, and that deletion of *SML1* derepresses RNR thereby promoting mtDNA synthesis. Indeed, we found that the steady state levels of Sml1 are drastically increased in cells disrupted of *IRC25* and *POC4* (see Fig. 4). The data strongly support the idea that mtDNA instability in the 20S proteasomal mutants is mediated by the stabilization of Sml1, which in turn inhibits RNR activity and dNTP biosynthesis.

**Figure 4.**
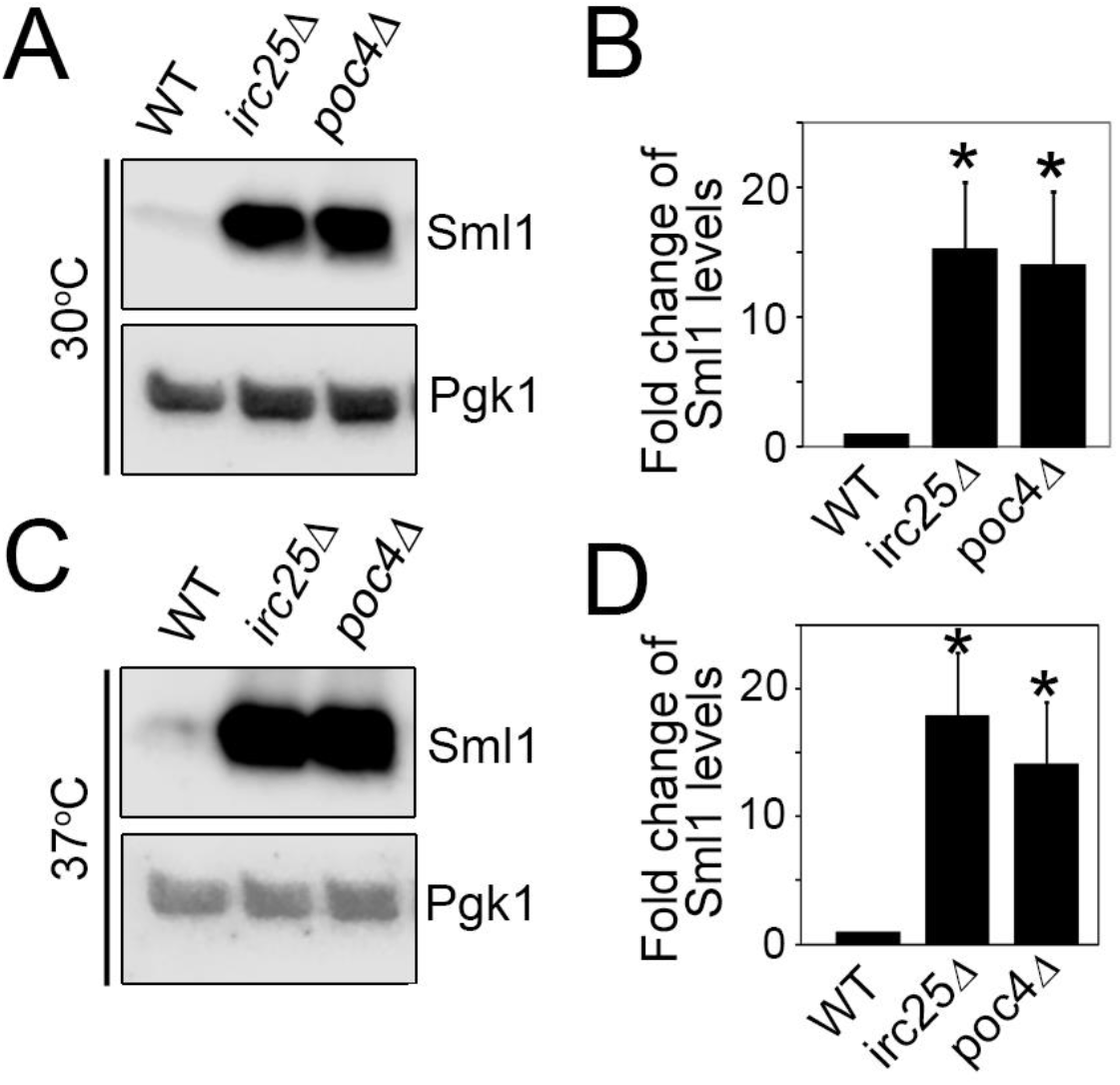
Stabilization of Sml1 in *irc25Δ* and *poc4Δ* mutants. (A) and (C) Western blot showing stabilization of Sml1 in *irc25Δ* and *poc4Δ* mutants grown at 30°C and 37°C. (B) and (D), quantitation of (A) and (C) respectively. *, *p* < 0.05.

### Loss of 19S proteasomes has subtle effect on Sml1 degradation and mtDNA stability

We then examined whether defect in the 19S regulatory complex of the proteasome affects mtDNA stability. Cells with disrupted *NAS2, NAS6, RPN14* and *TMA17*, encoding chaperones required for the assembly of the 19S regulatory particle of the proteasome (Roelofs *et al*. 2009; Saeki *et al*. 2009; Hanssum *et al*. 2014) are viable (Fig. 5A). Surprisingly, we found that disruption of these genes has little effect on mtDNA stability even when cells were grown at 37°C (Fig. 5B and 5C). It is possible that loss of these genes only partially impairs proteasomal function to an extent that is not sufficient to stabilise Sml1 and to affect mtDNA stability. To address this, we disrupted *HSM3*, encoding another 19S chaperone. The growth of *hsm3Δ* mutant cells is inhibited at 37°C (Fig. 5A), which suggests severe 19S proteasome defect. Interestingly, we found that the mutant cells only slightly increase petite production under these conditions. This is consistent with the observation that Sml1 is barely detectable in the 19S mutants including *hsm3Δ* (Fig. 5D and 5E). Taken together, these data suggest that the 19S regulatory particle is less critical relative to the 20S proteasome for Sml1 turnover and the maintenance of mtDNA.

**Figure 5.**
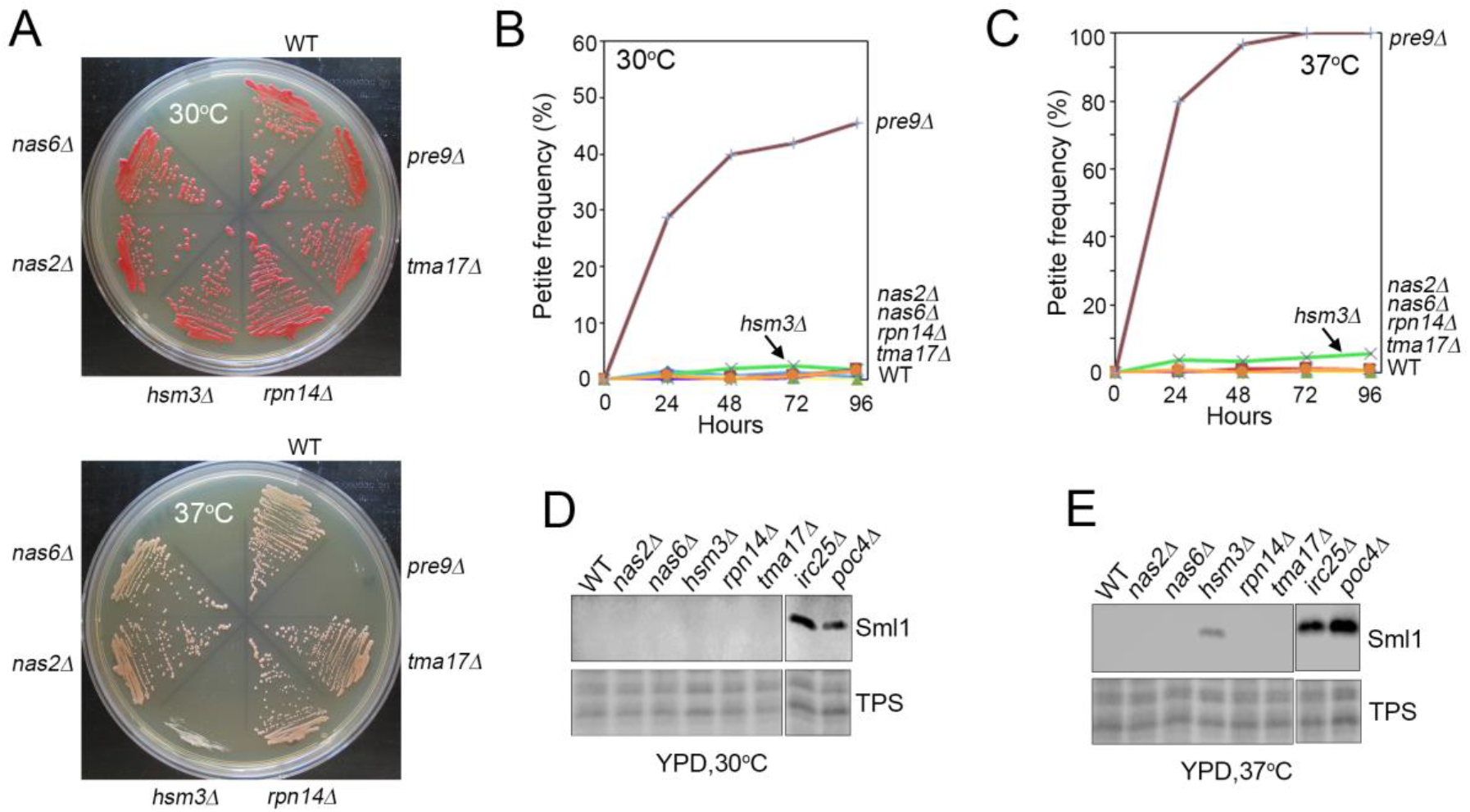
Effect of mutations in the 19S proteasome on mtDNA stability and Sml1 accumulation. (A) Growth of 19S proteasome mutants on YPD medium at 30°C and 37°C. (B) and (C) Petite frequency in 19S proteasome mutants grown at 30°C and 37°C respectively. (D) and (E) Sml1 levels in 19S proteasome mutants grown at 30°C and 37°C respectively. TPS, total protein staining.

### Sml1 level is decreased during oxidative growth and in cells with destabilized mtDNA

We hypothesized that Sml1-controlled dNTP synthesis may play a role in regulating mtDNA metabolism under conditions of mtDNA amplification and repair. In yeast, mitochondrial respiration is repressed by glucose during glycolytic growth and mtDNA copy number is increased when cells are shifted to oxidative growth. Indeed, we found that Sml1 levels reach the peak in exponentially growing cells (4 hours), begin to decline when cells enter into oxidative growth (8 hours), and become barely detectable entering into the stationary phase (Fig. 6A, upper panel). Although Irc25 and Poc4 are also decreased in oxidatively growing cells, the level of Pre9 is maintained (Fig. 6A, lower panel). This suggests the presence of an active 20S proteasome in these cells that contributes to Sml1 turnover. Sml1 is slightly stabilized in *irc25Δ* and *poc4 Δ* mutants in stationary phase (Fig. 6A, upper panel). We found that Sml1 is further stabilized during oxidative growth in *irc25Δ pep4Δ* and *poc4Δ pep4Δ* double mutants (Fig. 6B). Pep4 plays an important role in protein turnover in the vacuole. Our data indicate that both 20S proteasome and the vacuole participate in the degradation of Sml1 during oxidative growth, with the 20S proteasome playing a major role in the process.

**Figure 6.**
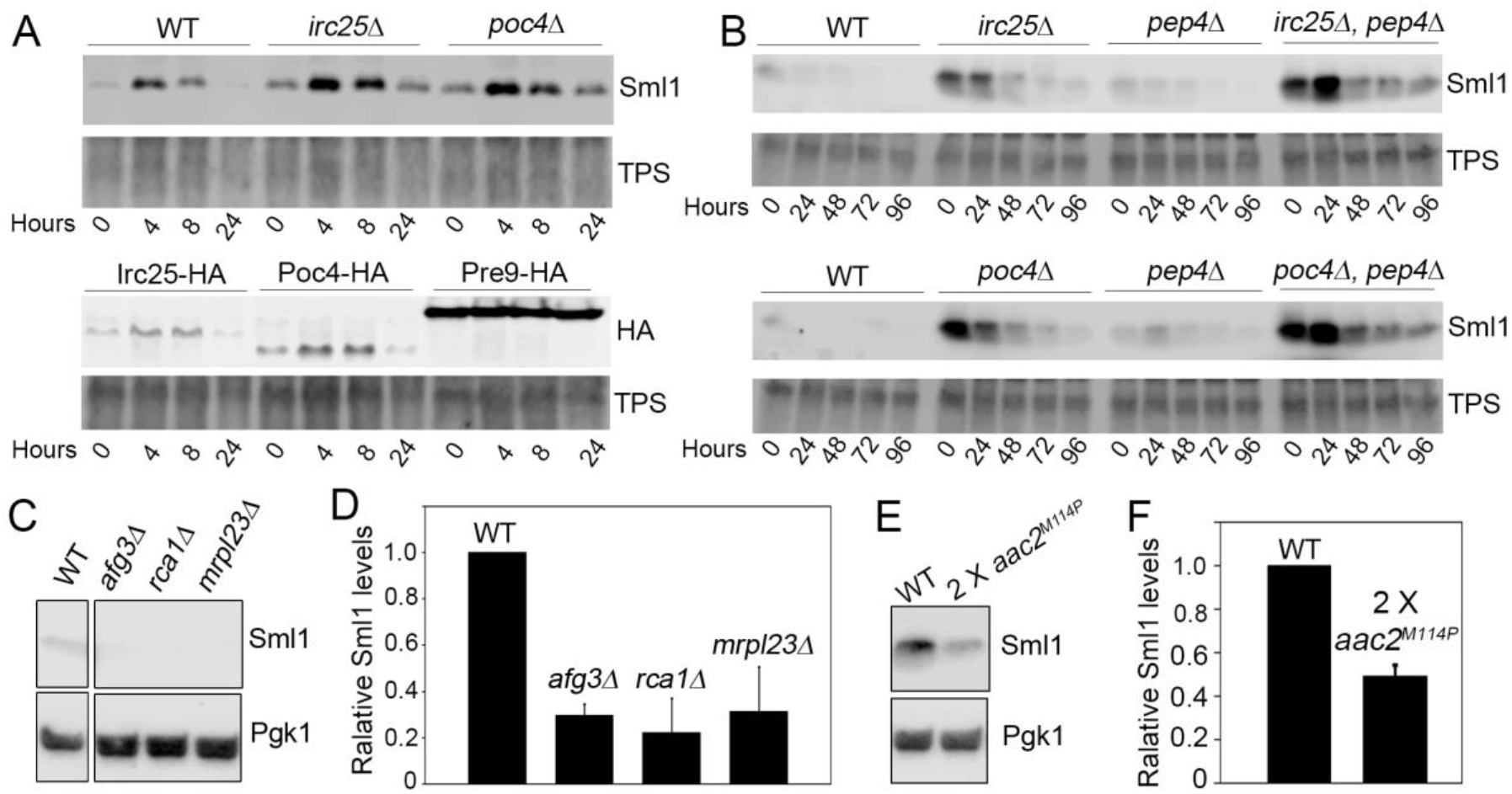
Sml1 levels at different metabolic states and in cells with destabilized mtDNA. (A) Accumulation of Sml1 (upper panel), Irc25-HA, Poc4-HA and Pre9-HA (lower panel) at different growth stages in YPD medium (30°C). TPS, total protein staining. (B) Western blot showing that degradation of Sml1 in stationary phase is dependent of 20S proteasome and vacuolar function. Cells were gown in YPD 30°C for 24, 48, 72 and 96 hours without refreshing medium. (C) Western blot showing reduced Sml1 accumulation in *afg3Δ, rcalΔ* and *mrpl23Δ* mutants. (D) Quantitation of (C). (E) Western blot showing reduced Sml1 accumulation in cells expressing 2 chromosomally integrated *aac2^M114P^*. (F) Quantitation of (E).

In addition to oxidative growth, we found that the steady state levels of Sml1 is significantly reduced in cells disrupted of *AFG3, RCA1* and *MRPL23* (Fig. 6C and 6D) and in a strain expressing two chromosomally integrated copies of *aac2^M114P^. AFG3* and *RCA1* encode two subunits of the i-AAA protease on the inner mitochondrial membrane. *MRPL23* encodes a subunit of the mitochondrial ribosome. mtDNA is highly unstable in these three mutants. mtDNA is also highly unstable in cells expressing *aac2^M114P^* (see Fig. 1A). These data support the idea that cells may respond to mtDNA instability by decreasing Sml1 levels as a compensatory mechanism to derepress RNR activity and to promote mtDNA repair and replication.

## DISCUSSION

The replication of nuclear and mitochondrial DNAs is not obligately coupled (Newlon and Fangman 1975; Sazer and Sherwood 1990). This is particularly important for quiescent cells in which mtDNA metabolism is required to maintain metabolic capacity in the absence of nuclear DNA replication and cell division. The replication and amplification of mtDNA is highly susceptible to dNTP level fluctuation in yeast (Taylor *et al*. 2005; Bradshaw *et al*. 2017; Puddu *et al*. 2019), supporting a critical role of the dNTP pools in regulating mtDNA metabolism. However, the bulk of cellular dNTP is synthesized during the S phase of cell cycle. How mtDNA metabolism is maintained in non-dividing cells is poorly understood. One possibility is that a rather basal level of dNTP synthesizing activity is maintained in the cytosol throughout the cell cycle. This may be necessary to meet the demands for mtDNA replication and repair, without interfering with cell cycle progression that may result from constitutively high levels of dNTP (Chabes and Stillman 2007). The Sml1 protein is an important inhibitor of RNR. Cells lacking Sml1 have higher levels of dNTPs, independent of increase in RNR transcription (Zhao *et al*. 1998; Tang *et al*. 2009). At S phase or after DNA damage, Sml1 is phosphorylated and degraded by the proteasome. This leads to the upregulation of dNTP by RNR that stimulates DNA synthesis and repair (Zhao and Rothstein 2002; Andreson *et al*. 2010). A previous study has observed mtDNA instability in the yeast checkpoint kinase Dun1 mutant that fails to eliminate Sml1 at S phase (Zhao and Rothstein 2002). In this report, we show that defect in the 20S proteasome can directly destabilize mtDNA due to Sml1 overaccumulation. The function of 20S proteasome is dynamically controlled under different physiological conditions. Out data support a 20S-dependent novel mechanism of RNR regulation and dNTP homeostasis that may be important for mtDNA maintenance and oxidative metabolism in non-dividing cells.

We found that yeast cells disrupted of *IRC25, POC4, BLM10, RPN4, UMP1* and *PRE9* have drastically destabilized mtDNA. *IRC25* and *POC4* are involved in the assembly of the 20S proteasome. *BLM10* is an activator of 20S (Schmidt *et al*. 2005). *RPN4* encodes a transcriptional factor of proteasomal genes. *PRE9* encodes the α3 subunit of 20S. These observations strongly support a role of the 20S proteasome in mtDNA maintenance. Multicopy suppressor screen revealed that overexpression of *RNR3* suppresses mtDNA loss from *poc4Δ* cells. More importantly, we found that mtDNA loss from the 20S mutants is almost completely suppressed by disruption of *SML1*. Previous studies have shown mtDNA instability in the *blm10* and *ump1* mutants, which was explained by impaired mitochondrial fission, increased production of reactive oxygen species and defective mismatch repair in mitochondria (Malc *et al*. 2009; Tar *et al*. 2014). We found that Sml1 is drastically stabilized in 20S mutants, which would be expected to suppress RNR activity. Based on these data, we conclude that mtDNA instability in the 20S mutants is mainly caused by reduced dNTP synthesis. Our finding provides an explanation for increased sensitivity of *blm10* mutants to genotoxins including hydroxyurea (Doherty *et al*. 2012). In all the 20S mutants examined with the exception of *rpn4Δ*, we found that incubation at 37°C accelerates mtDNA loss. It has been shown that *rpn4* mutant has reduced growth at normal temperature but is indistinguishable from the wild type cells at 37°C. It has been proposed that compensatory pathways may stimulate proteasomal genes independent of Rpn4 under the heat shock conditions (Xie and Varshavsky 2001; Hahn *et al*. 2006), which is likely sufficient to maintain 20S activity. We found that global proeostatic stress induced by the misfolded human Htt103Q protein also destabilizes mtDNA and this effect is fully suppressed by the disruption of *SML1*. Thus, general cellular proteostasis could also modulate dNTP homeostasis and mtDNA stability, possibly by affecting the availability of 20S proteasome for the degradation of Sml1.

It is interesting to observe that loss of 19S function seems to have very subtle effect on mtDNA stability. When the *hsm3Δ* mutant was incubated at non-permissive temperature to block 19S assembly, petite production was only marginally increased. Consistent with this, Sml1 accumulation is only slightly increased relative to what was observed in 20S mutants. This raises the possibility that the 20S proteasome may play an important role in the degradation of Sml1 independent of 19S. It has been extensively documented in the literature that the 20S proteasome, in contrast to the fully assembled 26S, is able to degrade unfolded proteins *in vitro* in an ATP- and ubiquitin-independent fashion (Davies 2001; Liu *et al*. 2003; Tsvetkov *et al*. 2009). This activity is often facilitated by specific proteasome activators specific to 20S (Opoku-Nsiah and Gestwicki 2018). It has been shown that shown that the N-terminal residues of α subunits play an important role in gating the proteolytic chamber of 20S (Bajorek *et al*. 2003; Choi *et al*. 2016), and that the substrate entry gate of 20S spontaneously opens to degrade peptides or unfolded proteins. In the absence of the α3 subunit, encoded by *PRE9*, yeast cells form the α4–α4 proteasome which has different catalytic activities. Loss of Irc25 and Poc4 required for α3 assembly would also be expected to favor the formation of α4–α4 proteasomes (Velichutina *et al*. 2004). Given that the *irc25* and *poc4* mutants cause overaccumulation of Sml1, the α3 subunit is therefore important for effective degradation of Sml1. It is known that Sml1 is ubiquitylated and its degradation is impaired in 20S mutants (Andreson *et al*. 2010). Future studies are required to learn whether non-ubiquitylated Sml1 is directly subjected to degradation by 20S independent of 19S and substrate ubiquitylation.

It is intriguing to observe that Sml1 is reduced to barely detectable levels when yeast cells enter stationary phase or quiescent state. This could be physiologically relevant. At the post-exponential growth phase, cell division is slowed down and oxidative metabolism is derepressed. Cells enter into a metabolic mode where mitochondrial mass is expanded and mtDNA copy number is increased. Interestingly, yeast cells at the stationary phase have been shown to increase proteasomal activity and to have increased release of 20S from the 26S proteasome (Fujimuro *et al*. 1998; Bajorek *et al*. 2003). We showed that the levels of 20S subunits such as Pre9 is maintained in the stationary phase. Both 20S activity and vacuole-mediated proteolytic activity are required for the degradation of Sml1 in stationary phase. Based on these data, we speculate that RNR activity is kept very low under normal growth conditions in non-S phase cells because of Sml1 inhibition. When transitioning to oxidative growth, increased degradation of Sml1 derepresses RNR and stimulates dNTP synthesis which in turn promotes mtDNA amplification and oxidative metabolism. Thus, the 20S proteasome serves as a novel regulatory loop to fine-tuning dNTP levels, allowing mtDNA replication when cells are switched to oxidative metabolism. This regulatory loop is vulnerable as demonstrated by the drastic destabilization of mtDNA in cells with 20S deficiency. Interestingly, we also observed that Sml1 is reduced in cells with destabilized mtDNA. It is known that mitochondrial damage activates proteasomal function in the cytosol (Wrobel *et al*. 2015) and is often marked by the release of 20S from the 26S proteasome (Livnat-Levanon *et al*. 2014). In this regard, it is possible that proteasome-based Sml1 degradation serves as a stress response that may retrogradely stimulate mtDNA metabolism by derepressing RNR activity.

Coupling dNTP homeostasis with proteostasis is not unprecedented. In *Caulobacter crescentus*, increased proteostatic stress reduces the availability of the Lon protease that normally degrades the transcriptional factor CcrM involved in dNTP production (Zeinert *et al*. 2020). As such, dNTP levels are increased to maintain replication capacity when midfolded proteins burden increases. In light of the current study, yeast cells appear to couple proteostasis with dNTP homeostasis to regulate mtDNA metabolism independent of cell cycle. In mammalian cells, the p53R2 subunit seems to play a key role in mtDNA maintenance (Pontarin *et al*. 2008). In light of the current study, it would be interesting to find out in the future whether activity of the R1/p53R2 enzyme is also regulated by the proteostatic status of the cell.

## ACKNOWLEDGEMENTS

We thanks Rodney Rothstein for providing the anti-Sml1 antibody and Brooke Hamling for help in yeast strain construction. This work was supported by the NIH grants AG061204 and AG063499 to X.J.C., and F30AG-060702 to L.P.C..

---

**Table 1.**
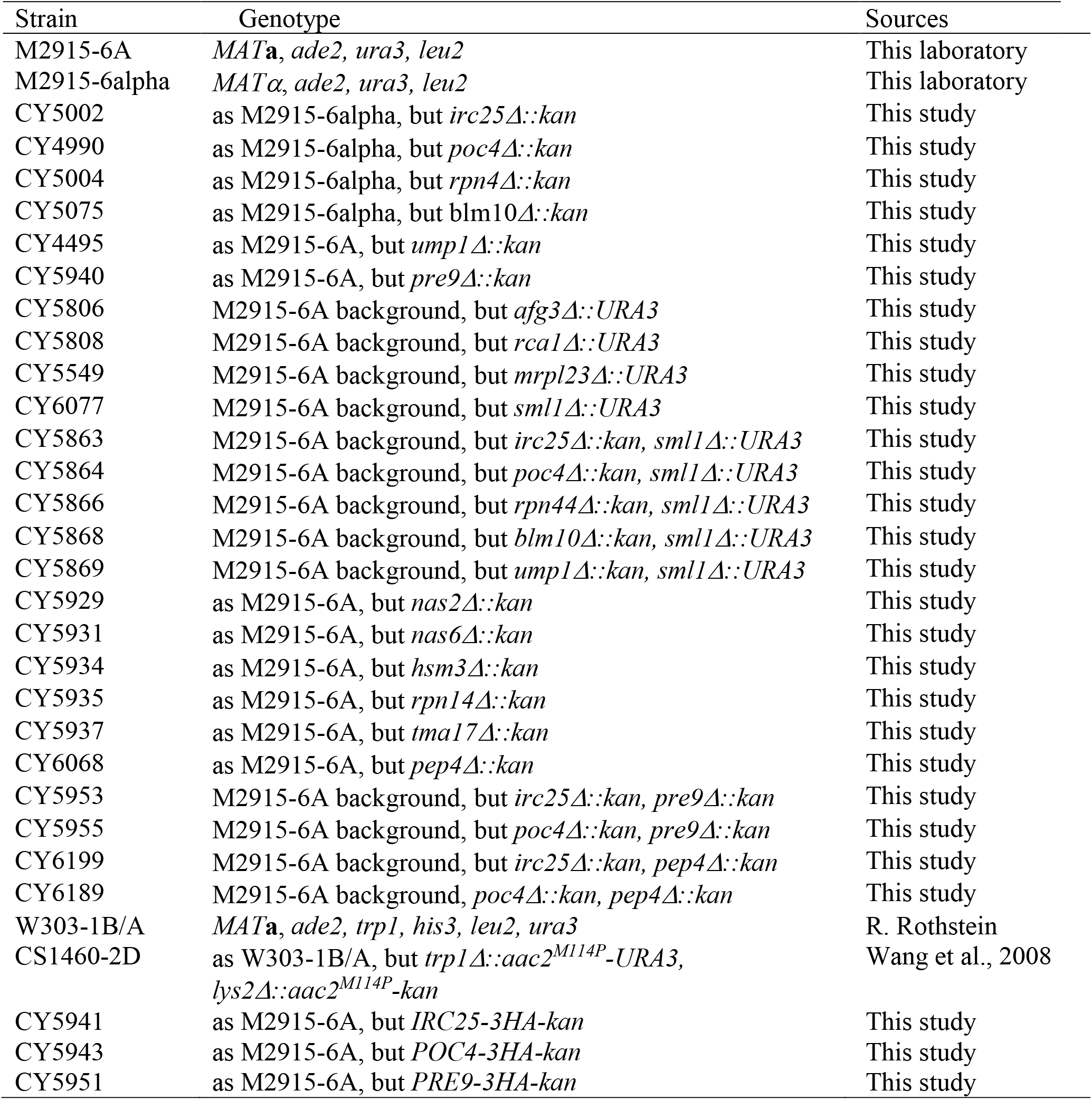
Yeast strains used in the study.

| Strain | Genotype | Sources |
| --- | --- | --- |
| M2915-6A | *MATa, ade2, ura3, leu2* | This laboratory |
| M2915-6alpha | *MATα, ade2, ura3, leu2* | This laboratory |
| CY5002 | as M2915-6alpha, but *irc25Δ::kan* | This study |
| CY4990 | as M2915-6alpha, but *poc4Δ::kan* | This study |
| CY5004 | as M2915-6alpha, but *rpn4Δ::kan* | This study |
| CY5075 | as M2915-6alpha, but *blm10Δ::kan* | This study |
| CY4495 | as M2915-6A, but *ump1Δ::kan* | This study |
| CY5940 | as M2915-6A, but *pre9Δ::kan* | This study |
| CY5806 | M2915-6A background, but *afg3Δ::URA3* | This study |
| CY5808 | M2915-6A background, but *rca1Δ::URA3* | This study |
| CY5549 | M2915-6A background, but *mrpl23Δ::URA3* | This study |
| CY6077 | M2915-6A background, but *sml1Δ::URA3* | This study |
| CY5863 | M2915-6A background, but *irc25Δ::kan, sml1Δ::URA3* | This study |
| CY5864 | M2915-6A background, but *poc4Δ::kan, sml1Δ::URA3* | This study |
| CY5866 | M2915-6A background, but *rpn44Δ::kan, sml1Δ::URA3* | This study |
| CY5868 | M2915-6A background, but *blm10Δ::kan, sml1Δ::URA3* | This study |
| CY5869 | M2915-6A background, but *ump1Δ::kan, sml1Δ::URA3* | This study |
| CY5929 | as M2915-6A, but *nas2Δ::kan* | This study |
| CY5931 | as M2915-6A, but *nas6Δ::kan* | This study |
| CY5934 | as M2915-6A, but *hsm3Δ::kan* | This study |
| CY5935 | as M2915-6A, but *rpn14Δ::kan* | This study |
| CY5937 | as M2915-6A, but *tma17Δ::kan* | This study |
| CY6068 | as M2915-6A, but *pep4Δ::kan* | This study |
| CY5953 | M2915-6A background, but *irc25Δ::kan, pre9Δ::kan* | This study |
| CY5955 | M2915-6A background, but *poc4Δ::kan, pre9Δ::kan* | This study |
| CY6199 | M2915-6A background, but *irc25Δ::kan, pep4Δ::kan* | This study |
| CY6189 | M2915-6A background, *poc4Δ::kan, pep4Δ::kan* | This study |
| W303-1B/A | *MATa, ade2, trp1, his3, leu2, ura3* | R. Rothstein |
| CS1460-2D | as W303-1B/A, but *trp1Δ::aac2<sup>M114P</sup>-URA3, lys2Δ::aac2<sup>M114P</sup>-kan* | Wang et al., 2008 |
| CY5941 | as M2915-6A, but *IRC25-3HA-kan* | This study |
| CY5943 | as M2915-6A, but *POC4-3HA-kan* | This study |
| CY5951 | as M2915-6A, but *PRE9-3HA-kan* | This study |

