## Supplemental Figure for "Mitochondrial DNA metabolism is coupled with 20S proteasome function via regulation of deoxyribonucleotide homeostasis in *Saccharomyces cerevisiae*"

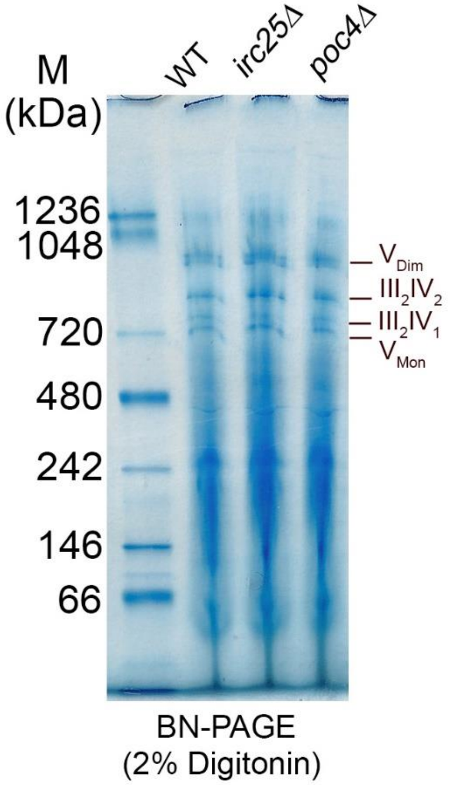


**Supplemental Figure 1. BN-PAGE analysis of respiratory complexes in *irc25Δ* and *poc4Δ* mutants compared with wild-type**. III, respiratory complex III; IV, respiratory complex IV; $V_{Mon}$, monomer of ATP $F_1F_o$-ATP synthase; $V_{Dim}$, dimer of $F_1F_o$-ATP synthase.
